# The fourspine stickleback (*Apeltes quadracus*) has an XY sex chromosome system with polymorphic inversions on both X and Y chromosomes

**DOI:** 10.1101/2024.10.21.619356

**Authors:** Zuyao Liu, Amy L. Herbert, Yingguang Frank Chan, Marek Kučka, David M. Kingsley, Catherine L. Peichel

## Abstract

Teleost fish are well-known for possessing a diversity of sex chromosomes and for undergoing frequent turnovers of these sex chromosomes. However, previous studies have mainly focused on variation between species, while comparatively little attention has been given to sex chromosome polymorphisms within species, which may capture early stages of sex chromosome changes. To better understand the evolution of sex chromosomes, we used the fourspine stickleback (*Apeltes quadracus*) as a model organism. Previously, it was believed that females of this species possessed a ZW heteromorphic sex chromosome system. However, genetic crosses and our whole-genome sequencing of wild populations revealed that *A. quadracus* has an XY sex chromosome on chromosome 23. This chromosome has not previously been identified as a sex chromosome in other stickleback species, indicating a recent sex chromosome turnover. We also identified two genes - *rxfp2a* and *zar1l* - as novel candidate sex determination genes. Notably, we observed inversions on both the X and Y chromosomes in different populations, resulting in distinctive patterns of differentiation between the X and Y chromosomes across populations. We propose that the inversion on the X chromosome may have been favored by sexually antagonistic selection. The new sex chromosome and intraspecies inversion polymorphisms observed in *A. quadracus* provide an excellent system for studying the evolution of sex chromosomes and their turnovers.

**Author Summary:** As compared to mammals and birds, teleost fish exhibit a very high level of diversity in their sex chromosomes, even among closely related species. Thus far, little attention has been paid to variation within species, although it offers a valuable opportunity to advance our understanding of the mechanisms underlying the formation and turnover of sex chromosomes. Through a quantitative trait locus (QTL) cross and sequencing diverse populations, we determined that instead of the previously reported ZW system, *A. quadracus* has an XY sex determination system on chromosome 23. Within the sex determining region, we identified *rxfp2a* and *zar1l* as putative sex determining genes. Notably, we also observed polymorphic inversions present on both the X and Y chromosomes that differ among populations. Based on our findings, we hypothesize that the X-linked inversions are favored by selection that differs between males and females. These observations represent a rare situation in which sex chromosomes are still polymorphic for sex-linked inversions, which offers important insights into the early stages of sex chromosome evolution.

## Introduction

Sex determination systems are diverse across species, and genetic sex determination mechanisms associated with the presence of heteromorphic sex chromosomes have independently evolved many times across the tree of life [1]. There are two main types of sex chromosomes. When males are the heterogametic sex, as in mammals, females have two X chromosomes and males have an X chromosome and a Y chromosome. When females are the heterogametic sex, as in birds, females carry a Z and a W chromosome, and males have two Z chromosomes. Although some groups like mammals and birds have very stable sex chromosome systems, in other groups like frogs [2,3], lizards [4], and fishes [5–7], even closely related species have different sex chromosome systems [8].

This diversity of sex chromosome systems is due to sex chromosome turnover, which occurs either when an existing sex determination gene moves to a new chromosome or when a novel sex determination gene arises on a chromosome [9,10]. According to the classical model of sex chromosome evolution, the acquisition of a new sex determination gene on an autosome can lead to the loss of recombination between the newly evolving proto-X and Y (or Z and W) chromosomes, resulting in the accumulation of deleterious mutations on the sex-specific and therefore non-recombining chromosome (Y or W) and the eventual formation of the heteromorphic sex chromosome pair [11,12]. This process can be interrupted by sex chromosome turnover, which resets the cycle and initiates the process again [9,10]. Although the evolutionary forces driving these turnovers are still unknown, sex chromosome turnovers have been hypothesized to occur due to selection for linkage between sexually antagonistic alleles and a sex-determination locus [13], selection to purge deleterious mutations that have accumulated on sex chromosomes[14,15], selection to maintain unbiased sex ratios [16,17], or random genetic drift [16,18,19]. However, testing these hypotheses remains challenging because it is difficult to catch sex chromosome turnovers when they are occurring [10].

The presence of polymorphic sex chromosomes within species would provide an opportunity to catch turnovers at an early stage. Intraspecies sex chromosome variation has been found in different groups, including frogs [2,20–22] and fishes [23–28]. Studies of these polymorphic systems have provided some insights into the evolutionary forces driving turnovers. For example, invasion of a new sex chromosome in cichlids is associated with linkage to a trait under sexually antagonistic selection [26]. Population-specific variation in the presence of a sex chromosome in a frog species is consistent with selection to purge deleterious mutations on the sex-specific chromosome [20]. Despite these limited examples, the evolutionary drivers and genetic mechanisms that underlie intraspecies variation in sex chromosomes are still mostly unknown.

A variety of sex chromosome systems have been identified in the species of the stickleback family (Gasterosteidae) that have diverged within the past 27 million years [29], suggesting that there have been recent sex chromosome turnovers. The three species in the genus *Gasterosteus* possesses a conserved heteromorphic sex XY sex chromosome on chromosome 19, with the anti-mullerian hormone Y (*amhy)* gene as the candidate sex determination gene [30,31]. The ancestral Y chromosome has independently fused to different autosomes in *G. nipponicus* and *G. wheatlandi* [7,32,33]. In a separate stickleback genus, independent duplication of *amh* has also been found as a candidate sex determination gene on chromosome 20 of *Culaea inconstans* [29]. Chromosome 12 is involved in an XY sex determination system in some *Pungitius* species, and another ZW sex determination system on chromosome 7 has been detected in *P. sinensis* based on genetic mapping [34–37]. Even so, sex chromosomes have not yet been fully identified in other species in Gasterosteidae, therefore, additional work is needed to explore the origins and evolution of sex chromosome evolution and turnovers in this family.

Fourspine sticklebacks (*Apeltes quadracus*) are of interest, as previous studies suggest they might possess a different sex chromosome than other stickleback species, but the sex chromosome has still not been identified. Initially, cytogenetic analysis of a population from Maine, USA indicated that *A. quadracus* has a heteromorphic ZW sex chromosome [38]. In the years following this study, conflicting evidence from new cytogenetic analyses were reported. Females from a Massachusetts (MA), USA population were found to have a heteromorphic sex chromosome, while no heteromorphic sex chromosome was identified in females (or males) from a Connecticut (CT), USA population [7,39]. These data suggested that the sex chromosome system might be polymorphic within *A. quadracus*, making this species an attractive target for further study of the evolution of sex chromosome turnover.

To identify the sex chromosomes in fourspine stickleback, we used several distinct approaches, including taking advantage of a published quantitative trait locus (QTL) cross done in wild populations of *A. quadracus* from Nova Scotia, Canada [40]. Additionally, we collected wild samples from three additional populations, two of which (MA and CT) were used for the previous cytogenetic studies, and one of which (a third population from Nova Scotia, Canada, NS hereafter) was used to generate a de novo genome assembly (S1 Fig). Then, for each population, we created crosses from a single mother and father per population and used pooled sequencing data (Pool-seq) from the crosses to identify the sex chromosome and sex determination region (SDR) in fourspine stickleback. For each population, we also generated haplotagging linked-reads sequencing data from wild individuals of both sexes. We further used these data to explore the variation on the sex chromosome among populations and identify candidate sex-determination genes.

## Results

### Genetic mapping of sex in a Nova Scotia (NS) intercross

Utilizing a previously described QTL cross between two *A. quadracus* populations from NS [40], we scored male vs. female status and mapped sex to chromosome 23 of the existing female assembly [41] (logarithm of odds (LOD) score 93.43, percent variance explained = 69.23%) (Fig 1A). Surprisingly, when we examined the genotypes of fish at the peak marker on chromosome 23, males overwhelmingly appeared to be the heterogametic sex, indicating that *A. quadracus* has an XY sex determination system (Fig 1B).

**Fig 1.**
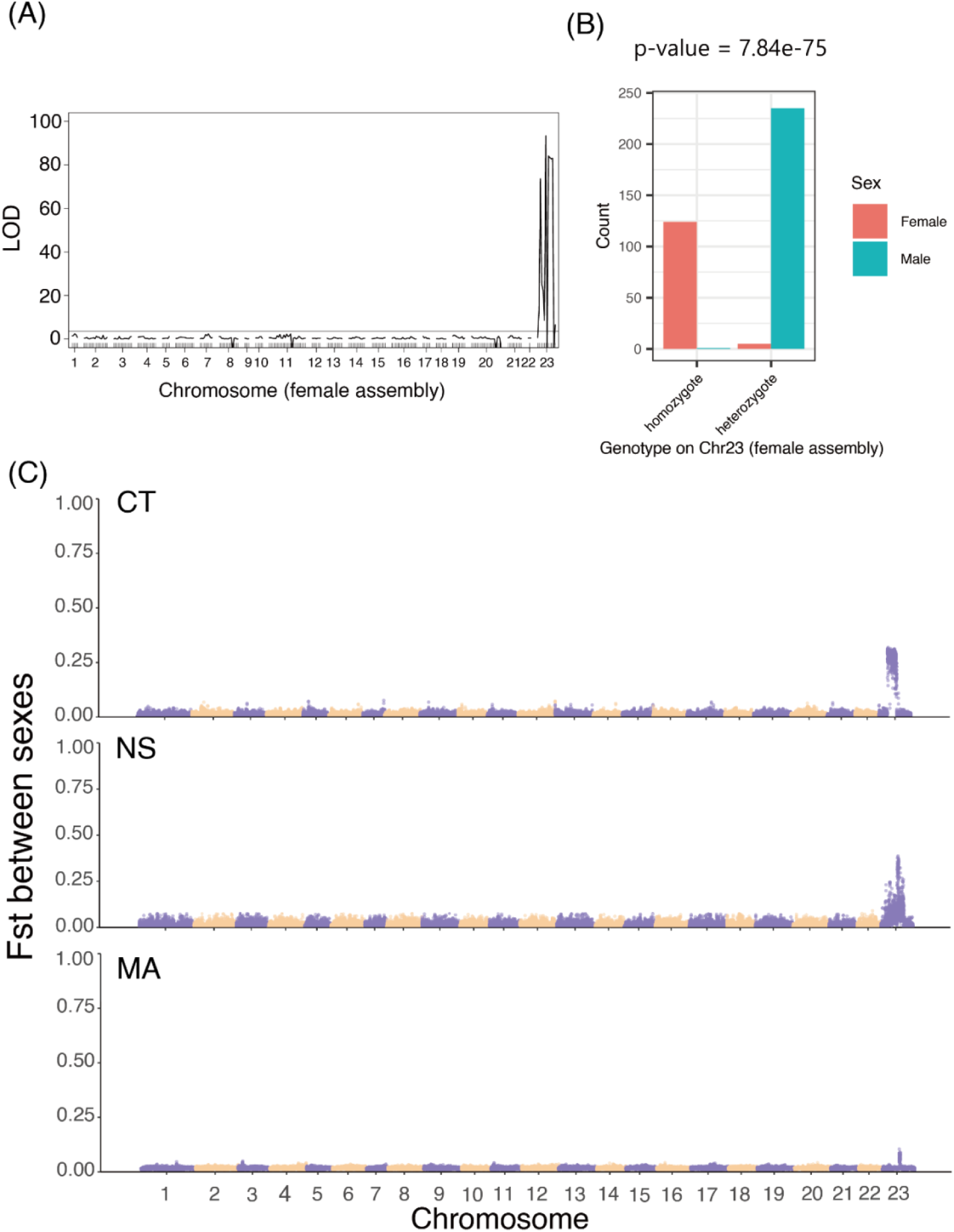
Identification of an XY sex chromosome system on chromosome 23. (A) QTL mapping of sex identifies a strong signal on *A. quadracus* chromosome 23 in the female genome assembly. The horizontal line shows the threshold for obtaining genome-wide significance with 1,000 permutations of the data and α = 0.05. (B) Bar plots show a highly significant correlation of genotype to phenotype at the top QTL peak marker, indicating that males are most likely the heterogametic sex. *P* = 7.84 e-75 (Chi-square test). (C) Genomic distribution of fixation index (Fst) between males and females in wild populations from Connecticut (CT), Nova Scotia (NS), and Massachusetts (MA). The size of the sliding window is 20 kb and the step size is 10kb. Chromosomes are indicated on the X-axis, and the Fst values are shown on the Y-axis. Purple and yellow regions indicate the different

### De novo assembly and annotation of a male *A. quadracus* genome

The existing *A. quadracus* genome assembly used in the QTL mapping analysis was generated from a NS female [41]. Due to the discovery of an XY sex determination system, we also generated a new high-quality assembly of a NS male genome from the same population using high-coverage PacBio HiFi and HiC reads. Raw HiFi read coverage was 116.79x (46.72 Gb in total) and HiC read coverage was 153.78x (63.05Gb in total). The final assembly is 475.93 Mb, and it contains 1261 scaffolds, including 24 chromosome-level scaffolds. The N50 length is 18.49 Mb, and the assembly quality assessed by BUSCO was relatively high with 97.4% completeness. We constructed a repeat library for *A. quadracus* using de novo-based approaches (see Materials and Methods). After masking the repetitive regions, the rest of the genome was annotated with evidence from brain RNA-seq data, homologous protein databases, and ab initio annotation, leading to 21,805 genes in the final version of the annotation. All analyses that follow are aligned to the X chromosome of this male reference genome; thus, all coordinates provided hereafter refer to the position on the X chromosome.

### Genome-wide analysis confirms an XY sex determination system on chromosome 23

To confirm our findings from the QTL mapping, we first utilized Pool-seq data from crosses of each of three populations (CT, NS, and MA in S1 Fig). Because these data are from siblings of relatively small genetic crosses (S1 Table), the number of recombination events limits our ability to define the non-recombining SDR. For example, any SNPs that are in the recombining region in the father may look sex-linked in the offspring if they did not happen to recombine with the X chromosome. Hence, we also conducted linked-reads sequencing of additional wild samples from the same three populations (20 females and 20 males from the CT population, 15 females and 14 males from the NS population, and 13 females and 11 males from the MA population).

The ratio of sequencing depth between males and females shows no discernible differences on any chromosome in either the crosses (S2 Fig) or the wild population linked-read sequences (S3 Fig), suggesting that heteromorphic sex chromosomes with large structural deletions are absent. However, if smaller mutations specific to a sex-specific chromosome have accumulated, increased genetic differentiation between the two sexes as well as increased diversity within the heterogametic sex on the sex chromosome would be expected. Consistent with the QTL mapping data, the fixation index (Fst) between males and females is elevated on chromosome 23 in both the crosses (S4 and S5 Figs) and the wild fish (Figs 1C and 2) from all three populations. Furthermore, diversity (Pi) is elevated in males, but not females, on chromosome 23 in the CT and NS crosses (S6 and S7 Figs) and the wild fish of all populations (Fig 2 and S8 Fig). Higher diversity in males is consistent with an analysis of RNA-seq data from the NS cross by SEX-DETector [42], which showed more sex-linked transcripts with evidence of male heterogamety than female heterogamety (S2 Table). Thus, there is evidence that all populations have a shared XY sex determination system on chromosome 23.

**Fig 2.**
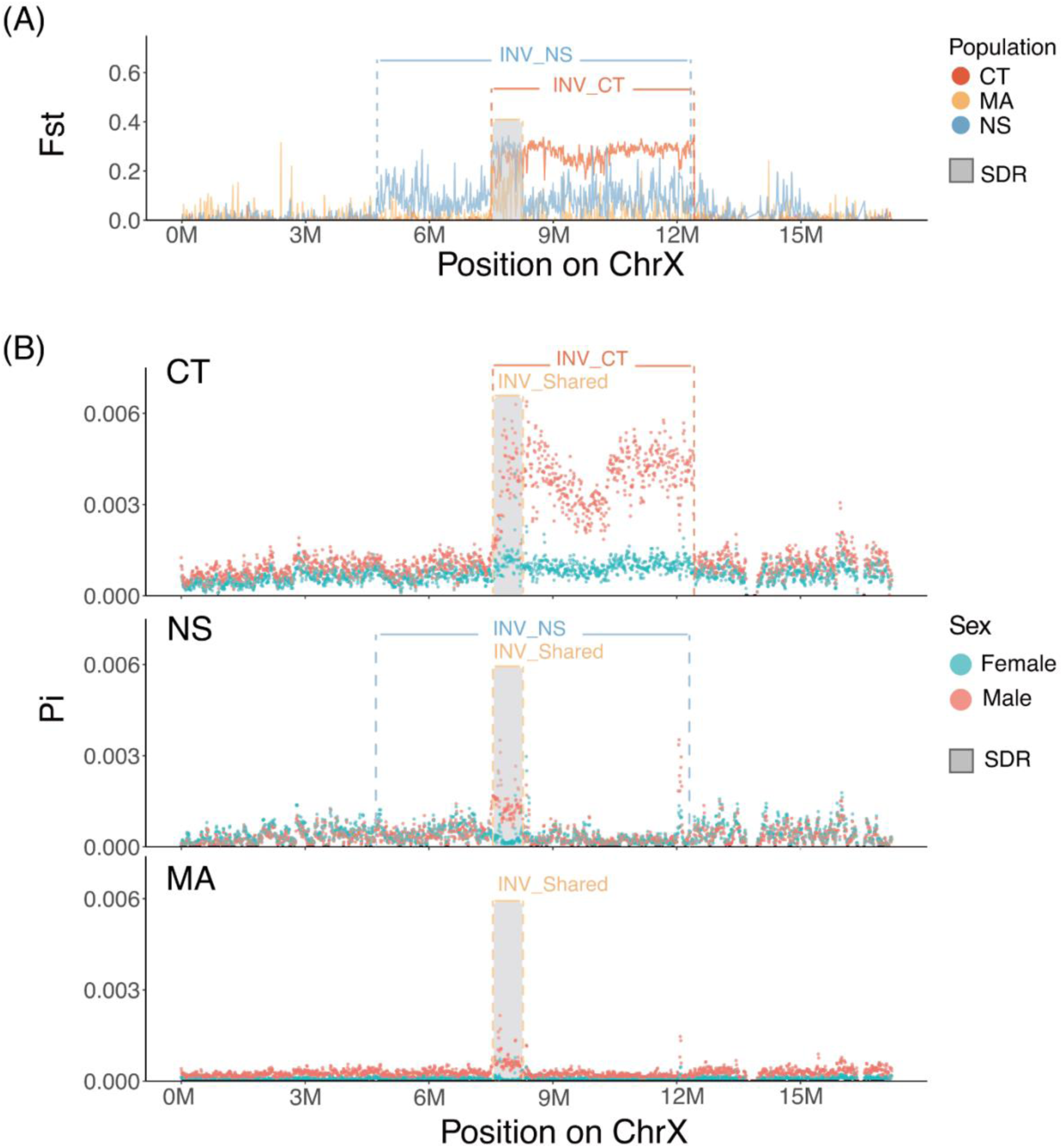
Patterns of genetic differentiation and diversity on chromosome 23 in wild populations. (A) Genetic differentiation (Fst) between males and females along chromosome 23 was calculated with linked-reads sequencing data from three wild populations, with the Connecticut (CT) population in coral color, the Massachusetts (MA) population in yellow, and the Nova Scotia (NS) population in light blue. The coral and light blue lines represent the corresponding inversions identified in the CT and NS populations, and the yellow lines represents a shared inversion among all populations. (B) Distribution of genetic diversity (Pi) with males (red dots) and females (cyan dots) on chromosome 23 calculated using linked-reads sequencing data from the three wild populations. The grey region represents the shared sex-determination region (SDR) across the three populations. Note that all sequences are aligned to the X chromosome assembly.

### Different populations have different X- and Y-linked inversions

Despite the presence of a shared XY sex determination system, the patterns of differentiation and diversity on chromosome 23 differed among the populations. In the CT cross, the differentiation between males and females (S5 Fig) and genetic diversity in males (S7 Fig) is elevated from 7.50 to 12.50 Mb on the X chromosome. Differentiation and genetic diversity show similar patterns across the X chromosome in the wild CT population (Fig 2): differentiation between males and females is elevated from 7.50 to 12.50 Mb, and males have higher diversity than females in this same region. The sharp boundaries of elevated divergence and diversity suggest that there is a rearrangement between the X and the Y chromosome. Indeed, analysis of the linked-reads from the CT wild population reveals that there is an inversion between 7.50 and 12.50 Mb that is polymorphic in both sexes (Fig 2). Specifically, 17 of 20 females are homozygous for the inverted orientation, while 17 of 20 males are heterozygous for the inverted orientation (S3 Table and S1 Appendix). Hence, we inferred that the inversion is on the X chromosome, which explains its existence in both sexes. The presence of this X-linked chromosome inversion explains the high genetic differentiation between males and females in both the cross (S5 Fig) and population data (Fig 2A). The lack of elevated diversity in females from the CT cross (S7 Fig) suggests that the mother of this cross was homozygous for the X-linked inversion and that the father also had an X chromosome with the inversion such that all daughters were homozygous for the inversion on the X chromosome and all sons are heterozygous for the inversion (S9 Fig).

In the NS cross, genetic differentiation between males and females is elevated between 2.73 and 17.20 Mb (S5 Fig); however, higher genetic diversity is observed only in males between 7.50 and 8.21 Mb (S7 Fig). The results of Fst and genetic diversity using linked reads from the wild NS population further refine the regions of elevated differentiation. There is a moderate elevation in genetic differentiation between 4.82 and 12.75 Mb, with a pronounced peak between 7.50 and 8.21 Mb (Fig 2A). Elevated male diversity in the NS wild population is confined to the 7.50 to 8.21 Mb region (Fig 2B). Analysis of the linked-reads further identifies an inversion between 4.72 and 12.44 Mb that is heterozygous and specific to males, indicating it is a Y-specific inversion (S3 Table). The presence of this inversion is further confirmed by comparing the assemblies of the X and Y chromosomes in an NS male (Fig 3A and B). Despite the presence of this large inversion on the Y chromosome, both the cross and wild population data show that male genetic diversity is concentrated within the region between 7.50 and 8.21 Mb, suggesting limited divergence between the X and Y chromosomes in the inverted region (Fig 2B and S7 Fig). The region of elevated diversity around 12 Mb (Fig 2B), located near the breakpoint of the Y-specific inversion, may reflect the presence of repetitive elements rather than true differentiation between the X and Y chromosomes, as diversity is increased in both sexes.

**Fig 3.**
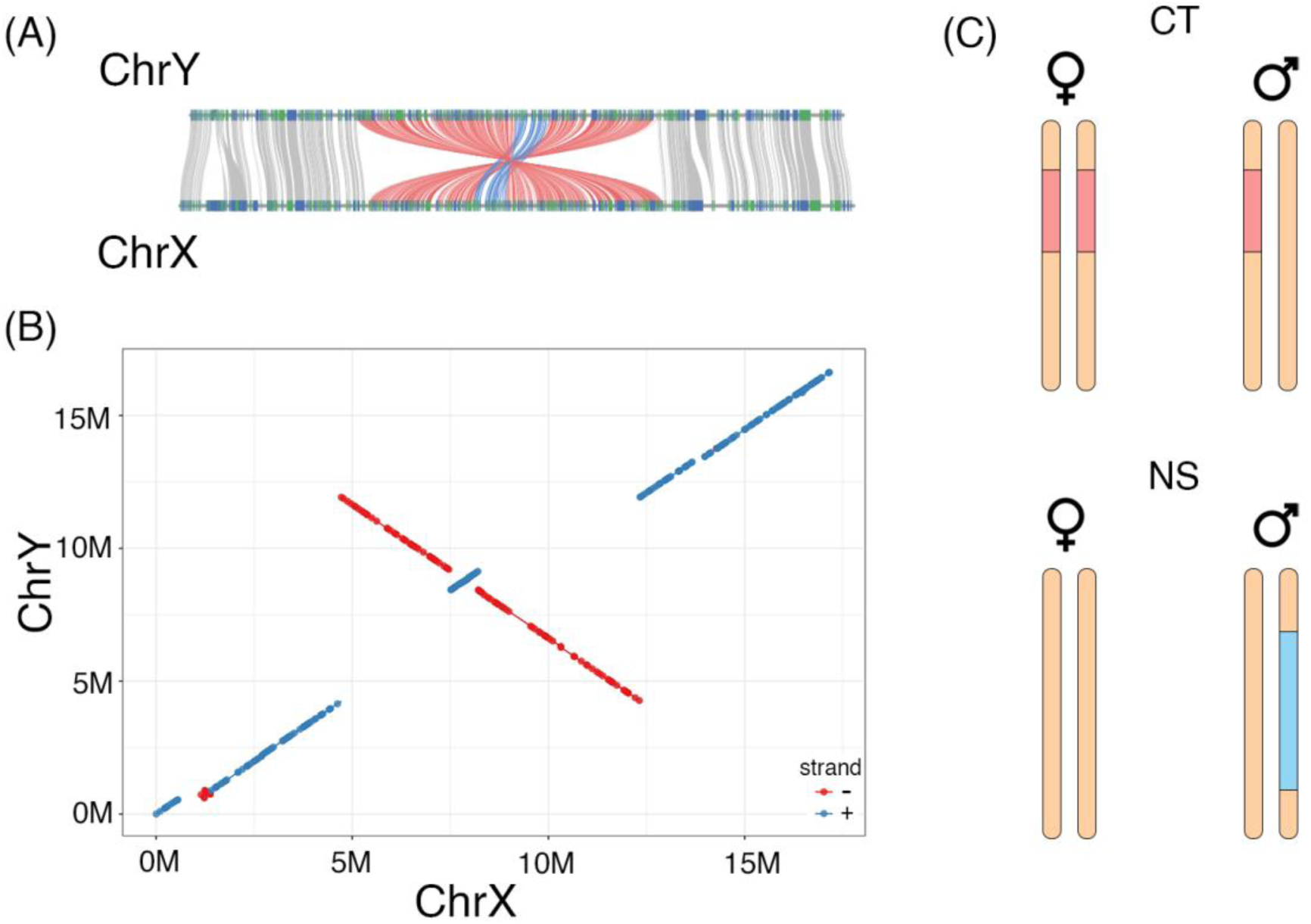
Shared and population-specific inversions on the X and Y chromosomes. (A) Synteny map between X chromosome (ChrX) and Y chromosome (ChrY) from the *A. quadracus* NS male assembly. This comparison is based on homologous coding region sequences. Colored lines are gene pairs. Red lines represent the larger inversion on the Y chromosome, and blue lines represent the nested inversion that covers the SDR. (B) Synteny map between ChrX and ChrY from the *A. quadracus* male assembly. This comparison is based on full sequences. Blue dots represent forward alignments, and red dots represent reverse alignments. (C) Model for population-specific inversions on the *A. quadracus* sex chromosomes. Orange bars represent the sex chromosome pair on chromosome 23. Coral bars show positions of the X-specific inversion in both sexes in the CT population, and the light blue bar shows the position of the Y-specific inversion in males in the NS population.

In the MA cross, genetic differentiation is elevated between 2.73 and 7.50 Mb, as well as from 12.50 to 17.20 Mb, with both sexes displaying high levels of genetic diversity within these regions (S5 and S7 Figs). However, in the linked-reads data from the wild MA individuals, Fst is primarily elevated between 7.50 and 8.21 Mb, and this same region exhibits an enrichment of genetic diversity specifically in males in the MA population (Fig 2). Direct assessment of potential rearrangements was challenging due to the insufficient average sequencing depth in the MA wild population, preventing confident genotyping of the inversions. Nevertheless, in the region associated with the X-linked inversion in the CT population, low divergence between males and females, coupled with high genetic diversity in both sexes, was observed in the cross data (S5 and S7 Figs). This pattern could be explained if the mother in the cross was heterozygous for the inversion, while the father lacked the inversion. In such a scenario, the inversion would be inherited equally by sons and daughters, resulting in no differentiation between the sexes and similar levels of genetic diversity in both (S9 Fig). These findings suggest that the X-linked inversion may be present at a low frequency within the MA population.

Summarizing the above evidence, we propose a model for the pattern of inversions in different populations (Fig 3C). The CT population has an X-specific inversion, whereas the NS population has a Y-specific inversion covering most of the Y chromosome. The two identified inversions in the CT and NS populations are derived, as they are inverted relative to the X chromosome assembly from the NS population, whose orientation appears to be ancestral by comparison to the genome assemblies of other sticklebacks (*G. aculeatus* and *P. pungitius*) and an outgroup species (*Aulorhynchus flavidus*) [41] (S10 Fig).

### Defining the shared sex determination region (SDR)

To define the shared SDR across the three populations, we focused on the linked-read data from the wild populations. A common region with elevated Fst is observed across all three populations, where higher genetic diversity is evident in males (between 7.20 - 8.21Mb on the X chromosome, Fig 2A). Additionally, the homologous region on the Y chromosome is enriched in male-specific kmers in all three populations, suggesting that they have a shared region with Y-specific alleles (S11 Fig). Furthermore, split reads and discordantly mapped read pairs suggest that there is a shared Y-specific inversion between 7.50 and 8.21 Mb (S3 Table). Due to the poor mapping quality, the genotype of individuals cannot be determined for this inversion. However, the synteny map between the X and Y chromosome assemblies in the NS population suggests that this inversion exists and is nested within the larger inversion on the Y chromosome (Fig 3A and B). Thus, we identify a shared SDR between 7.50 Mb and 8.21 Mb, which likely contains the primary sex determination gene. As this region is covered by the inversion shared among populations, the SDR might have been formed by an inversion.

### Lack of degeneration on the *A. quadracus* Y chromosome

Although the homomorphic sex chromosomes of *A. quadracus* show no large-scale regions of read depth reduction (S2 and S3 Figs), we explored whether there has been degeneration at a fine scale. One method to identify degenerated genes on the sex chromosomes involves comparing read depth of genes between males and females in wild populations using linked-reads data. If the ratio of male to female depth is less than 0.75, the gene is considered to be degenerate, indicating a loss of its content [33]. There are no genes on the X chromosome that are degenerate based on this criterion in either the CT or NS population (the MA population was excluded from these analyses given the low sequence coverage). We also looked for the presence of fixed loss-of-function mutations as evidence for degeneration. There are three loss-of-function mutations on the Y chromosome and two loss-of-function mutations on the X chromosome in the CT population, and three loss-of-function mutations on the Y chromosome and no loss-of-function mutations on the X chromosome in the NS population (S4 Table). As the differences are not pronounced, extensive degeneration has not occurred on the Y chromosome of *A. quadracus*.

### *Rxfp2a* and *zar1l* are candidate sex-determination genes in *A. quadracus*

Following the identification of the shared SDR, we conducted a thorough analysis of the genes within this region to identify potential candidates for primary sex determination. Our search for Y chromosome-specific genes revealed 8 predicted genes, of which four are in the SDR. Only three of the 8 genes had identifiable homologues based on BLAST analysis. Additionally, 34 genes were found to be present within the SDR on both the X and Y chromosomes. However, neither the four Y-specific genes nor the 34 genes shared between the X and Y chromosomes in the SDR show homology to known sex determination genes in other teleost species (S5 Table). There is also no homology between known sex determination genes and the five annotated genes present in the small region with elevated genetic diversity between 12.04Mb and 12.17Mb (Fig 2 and S5 Table). Thus, we focused our analyses on the genes present on the X and the Y within the SDR; for each gene, we counted the number of synonymous and nonsynonymous changes, separately for both sexes. Because this is an XY system, we focused on genes with changes between the X and Y chromosome. In total, there are 23 candidate sex-determination genes with nonsynonymous changes located on chromosome 23 within the SDR (S5 Table). Among these genes, there are two genes of interest, *rxfp2a* (7.565 Mb – 7.592 Mb) and *zar1l* (7.515 Mb – 7.516 Mb), which are related to the development of the reproductive system (see Discussion for details). There are three amino acid changes in the Y chromosome allele of *zar1l*, and all of them are predicted to cause a deleterious mutation, according to SIFT analysis [43]. The dN/dS ratio is 0.289. For the *rxfp2*a gene, the dN/dS ratio cannot be calculated as there are no synonymous mutations. There are two nonsynonymous mutations located in the coding region, but they are not predicted to have a significant effect on the function of the gene.

## Discussion

### Variation and turnover of sex chromosomes in *A. quadracus*

Previous evidence from cytogenetic studies suggested that populations of *A. quadracus* from Maine and Massachusetts have a heteromorphic ZW sex chromosome in females [7,38]. However, no heteromorphic sex chromosome was detected in metaphase spreads of males or females from Connecticut [39]. Using data from four genetic crosses and wild fish from three populations, we determined that *A. quadracus* has an XY sex determination system on chromosome 23. Our analyses included samples from both the Massachusetts and Connecticut populations used in the previous cytogenetic studies. Thus, the discovery that *A. quadracus* has an XY sex determination system is surprising, as the morphology of chromosomes in the MA population clearly indicated the presence of a heteromorphic pair in females [7]. One possible explanation for this result is if the MA individuals used for cytogenetics were heterozygous for the X-linked inversion that we identified in the CT population. If the inversion caused a change in chromosome morphology at the cytogenetic level, the chromosome pair may have appeared to be heteromorphic. Although we performed linked-reads sequencing of some of the MA females used for the cytogenetic study [7], we did not obtain high enough sequencing coverage to confidently assess inversion genotypes in these individuals. However, the MA cross data suggest that the X-linked inversion is present in the MA population (S5 and S7 Figs). As we do not have samples from the Maine population used in the older cytogenetic study [38], we could not assess whether the X-linked inversion is present in this population. If heteromorphic chromosomes in females are indeed due to heterozygosity for the X-linked inversion, it is not surprising that the CT females were homomorphic in the previous cytogenetic study since these females are mostly fixed for the inversion (S3 Table). However, to fully resolve this mystery, a more detailed molecular cytogenetic analyses of these different populations is needed, which will be facilitated by our identification of the SDR on chromosome 23 in *A. quadracus*.

Chromosome 23 has not previously been identified as a sex chromosome in sticklebacks, suggesting that there has been a sex chromosome turnover in *A. quadracus*. However, it is interesting to note that *A. quadracus* chromosome 23 is homologous to part of chromosome 7 in both *G. aculeatus* and *P. pungitius* [41]. The non-homologous part of chromosome 7 has fused to chromosome 12 in the *Pungitius* lineage, and there is evidence that chromosome 7 carries a female heterogametic (ZW) sex determination locus in *P. sinensis* and that chromosome 12 carries a male heterogametic sex (XY) determination locus in *P. pungitius* [34]. Given that the SDR in these two species is not homologous to that in *A. quadracus*, it is unlikely that they have the same sex determination gene. However, testing this hypothesis requires identifying the sex-determination gene in all three species. It is clear that *A. quadracus* has a different sex chromosome and sex determination gene from the *Gasterosteus* species, in which the master sex determination gene *amhy* is found on chromosome 19 [30,31,33,44], or in *C. inconstans*, in which there has been an independent duplication of *amhy* on chromosome 20 [29]. No duplicated copy of *amh* has been found in *Pungitius* species or in *A. quadracus* [29]. Further supporting a sex chromosome turnover in *A. quadracus* is the lack of extensive differentiation between the X and the Y or degeneration on the Y chromosome. Similar patterns on sex chromosomes in *P. pungitius*, *P. sinensis*, and *C. inconstans* hint that these turnovers also occurred quite recently [29,34,37], with evidence for a very recent additional turnover within *P. pungitius* [45]. The sex chromosomes in these species are in contrast to the Y chromosome in the *Gasterosteus* lineage, which evolved approximately 22 million years ago and has experienced extensive degeneration, albeit at different rates in the three species in this genus [30,31,33]. This variation in turnover among different stickleback lineages provides an opportunity to further investigate the factors that lead to sex chromosome stability in some lineages and turnover in others.

### Two novel candidate sex determination genes

We identified two novel candidate sex determination genes in the shared SDR on chromosome 23. Although we identified four Y-specific genes within the SDR, neither is a good candidate sex determination gene as none are homologous to the known sex-determining genes in teleosts (S5 Table). The genes *zar1l* and *rxfp2a* are the only two genes within the SDR that have Y-specific SNPs across all populations studied and are known to play roles in the development of the reproductive system (S5 Table).

The *rxfp2a* (relaxin/insulin-like family peptide receptor 2) gene encodes a receptor that plays a crucial role in the development of placental mammals by binding with high affinity to the peptide *INSL3* (insulin-like 3). This *INSL3/RXFP2* pairing is essential for the proper descent of the testicles during development in mammals [46], and loss of *rxfp2a* results in cryptorchidism in mice [47–50]. Phylogenetic analysis of 71 mammalian genomes revealed that the *rxfp2a* gene is lost or non-functional in four afrotherian species that lack testicular descent [51]. In zebrafish, *INSL3* regulates spermatogonia stem cell differentiation from mitosis to meiosis [52]. Since *rxfp2a* is a receptor for *INSL3*, mutations in this gene have the potential to disrupt the entire *INSL3/RXFP2* signaling pathway, ultimately affecting spermatogenesis. In *A. quadracus*, *rxfp2a* exhibits Y-specific mutations in both the coding and regulatory regions. Given the conserved role of this gene in testes development and spermatogenesis, *rxfp2a* is a candidate gene that warrants further investigation.

As a maternal effect gene conserved across vertebrates*, zar1* plays an important role in oocyte-embryo transition and impacts female fertility in mice [53]. In *Xenopus laevis*, the *zar1* gene controls the translation of Wee1 and Mos mRNAs in immature oocytes [54]. Additional evidence about *zar1* impacting the sex ratio was found in *Danio rerio*, where a complete male-biased sex ratio was observed in *zar1* knock-out mutants [55]. In addition, it was reported to have an effect on a number of known translation factors, such as CEPB, ePAB, and 4E-T [55,56], among which CPEB and ePAB are known for controlling the process of oogenesis [57] and 4E-T is associated with human primary ovarian insufficiency [47]. In our study, we found that there are two *zar1* genes in the *A. quadracus* genome: the ancestral copy is on Chr8, and the duplicated copy (*zar1l*) is found on both the X and Y copies of chromosome 23. The Y chromosome allele has three amino acid changes that are predicted to be deleterious. Considering this gene is quite conserved across species, it is likely that the amino acid changes disrupt the function. Hence, having one functional copy of *zar1* could lead to male development, which would be consistent with the zebrafish data. Therefore, we conclude that *zar1l* is another appropriate candidate gene. However, further experiments, such as gene knockouts and/or SNP editing by CRISPR-Cas9 are necessary to determine whether *rxfp2a* or *zar1l* is the master sex determination gene in *A. quadracus*.

While numerous sex determination genes have been identified and studied in fish, the two genes mentioned above, *rxfp2a* and *zar1l*, have not been previously identified as sex determination genes. In contrast, other genes, such as *amh*, *amhr2*, *dmrt1*, and *gdf6*, have been repeatedly identified as master sex determination genes in various fish species [58,59]. In stickleback species, one key sex determination gene is the independent duplication of the *amh* gene on the Y chromosome of *Gasterosteus* species [31] and *C. inconstans* [29]. However, there is no evidence for an additional copy of the *amh* gene on Chr23 (this study) or elsewhere in the *A. quadracus* genome [29], suggesting that *A. quadracus* has probably undergone a turnover in the sex determination gene.

Interestingly, some of the other genes that we highlight in the SDR on Chr23 (S5 Table) and two genes slightly outside this region have previously been shown to be linked to the SDR in a distant relative, the Atlantic herring (*Clupea harengus*). The sex determining gene in this group is likely *BMPR1BBY* [60] and the nearby linked genes include *mettl27*, *cldn4*, *smyd4*, *sms*, and *pgrmc1* [60,61]*. Smyd4* has high expression in zebrafish testis and *pgrmc1* has a role in zebrafish oocyte maturation, while the gene *sms* is linked to the X-chromosome in humans [61]. Both *mettl27* and *cldn4* are part of the cohort of genes deleted in the human disorder Williams-Beuren syndrome, which has been shown to affect genital development, with humans exhibiting phenotypes such as undescended testis, retractile testis, and cryptorchidism [62–64]. Although not thought to be primary sex determining genes in Atlantic herring, both *mettl27* and *cldn4* have SNPs that differ in males and females and lead to nonsynonymous amino acid changes [61]. This may therefore be a good example of genes with different functions in males and females becoming linked to the SDR. Although Atlantic herring and *A. quadracus* are diverged by approximately ∼220 million years of evolution [65], the finding of similar sex-linked genes in the two groups highlights the critical role that conserved supporting genes may play in sexual development and sex chromosome evolution.

### Polymorphic X- and Y-linked inversions on the sex chromosomes of *A. quadracus*

We have also identified polymorphic and derived inversions on both the X and Y chromosomes in *A. quadracus* populations. There was a high frequency of an X-linked inversion in both males and females in the CT population, partially covering the SDR (Fig 2A, S3 Table and S1 Appendix). Evidence for a similar X-linked inversion was also found with the Pool-seq data from the MA cross, indicating that it might be present at a low frequency in this population (S5 and S7 Figs). Although this X-linked inversion does not seem to be present in the NS population, discordantly mapped reads and shared barcodes point to a Y-specific inversion in this population (Fig 3 and S3 Table). Both the X and the Y-linked inversion contain the shared SDR from 7.50 Mb and 8.21 Mb, which coincides with a potential inversion that is shared among populations (Figs 2 and 3).

Inversions have been proposed as a mechanism to suppress recombination between X and Y chromosomes [66]. Several studies have now found evidence for Y-linked inversions associated with suppression of recombination on Y chromosomes [31,67–69]. A number of hypotheses have been proposed to explain the suppression of recombination on sex chromosomes, including sexual antagonism [12,70–73], meiotic drive [74], dosage compensation [75], sheltering of recessive deleterious mutations in heterozygotes [76,77], and genetic drift [78–80]. Our finding of a Y-linked inversion in the NS population is consistent with all of these models for suppression of recombination between the X and the Y. The case presented here could present a valuable opportunity to empirically test one or more of these theories. As the Y-linked inversion is polymorphic among populations, it allows us to observe the gene content and gene expression of both the ancestral and inverted Y haplotypes, as well as potentially comparing fitness of males with and without the inversion. We may also further sample the range of *A. quadracus* in search of populations for which the inversion is polymorphic. Such studies could shed light on the potential targets of selection within the inverted Y haplotype, or the lack thereof.

We also document here a polymorphic X-linked inversion, which is absent in the NS population and almost fixed in the CT population (and of unknown frequency in the MA population). The natural question to ask is whether this inversion is playing some role in the CT (and possibly MA) populations. Although a recent study in *Silene latifolia* also points to the presence of an X-linked inversion leading to suppression of recombination between X and Y chromosomes [81], there are currently no models which explicitly address the suppression of recombination on sex chromosomes via X-linked inversion. Furthermore, some of the models described above to explain the spread of Y-linked inversions cannot be easily applied to the X. The principal difference is that on Y chromosomes, inversions are specific to the heterogametic sex, and are therefore instantly and permanently heterozygous, which gives rise to their theorized fitness benefits (e.g. the sheltering of recessive deleterious alleles). In contrast inversions on the X can be present in both sexes and can be homozygous in females. This means that genes within an X-linked inversion cannot be sex-specific, and recessive deleterious alleles can only be sheltered by obligate heterozygosity in males, but not females, which would counter the spread of such inversions [77]. The meiotic drive model [74] is also unlikely to explain the presence of an X-linked inversion, as meiotic drivers often carry fitness costs in homozygous form [82], resulting in the exposure of deleterious alleles in females. The dosage compensation model [75] is also unlikely to explain the initial spread of an X-linked inversion because the inversion in the heterozygous state would lead to the incompatibility of expression modifiers in females. However, once fixed (regardless of why), the X-linked inversion could be maintained as the incompatibility of expression modifiers would prevent the restoration of recombination between the X and the Y. It is possible that drift could also play a role in the spread of X-linked inversions because recessive beneficial mutations, including inversions, do have a higher probability of fixation via genetic drift on X chromosomes than on autosomes. However, this is only the case when X-linked alleles are hemizygous in males, as in highly degenerate sex chromosomes [78]. As the *A. quadracus* Y chromosome has not experienced much degeneration, drift alone is not likely to be the explanation for the spread of the X-linked inversion.

Although more work is needed, we may still speculate on the circumstances leading to the spread and possible fixation of such inversions, based on the principles of X inheritance. Firstly, X-linked inversions are likely to be heterozygous more often than autosomal inversions, as homozygosity in males will not exist. Thus, theoretically a locus within the inverted haplotype which confers a fitness advantage to heterozygotes could favor the persistence and spread of an X-linked inversion under some circumstances. Secondly, as the X spends three-quarters of its time in females (as opposed to half of its time as in autosomes), there is scope for selection to favor alleles which benefit females, particularly if they are also deleterious in males (i.e., sexually antagonistic loci) [78]. Direct comparisons between the rate of fixation of X and Y-linked inversions under the sexually antagonistic selection hypothesis have not been done, but X-autosome fusions (which also could suppress recombination between the sex determination locus and a sexually antagonistic allele) can spread under sexually antagonistic selection, albeit more slowly than a Y-autosome fusion [71]. Consistent with the predictions of the sexual antagonism hypothesis, the inversion on the X chromosome does contain the shared sex determination gene. However, additional work is necessary to test this hypothesis, including assessing the frequencies of the X-linked inversion across many *A. quadracus* populations, and determining whether phenotypes under sexually antagonistic selection are associated with the inversion.

## Conclusions

Although variation in sex chromosomes systems among closely related species is now well-documented, the mechanisms behind sex chromosome turnover remain unclear. By examining population data from wild-caught samples and genetic crosses, we find evidence of a recent turnover in both the sex determination gene and the sex chromosome in *A. quadracus*. Furthermore, there are polymorphic inversions on the X and Y chromosomes, with relatively little degeneration on the Y chromosomes. This within-species variation on the *A. quadracus* sex chromosomes provides an opportunity for further studies to test hypotheses of the evolutionary forces driving sex chromosome evolution and turnover.

## Materials and Methods

### Ethics statement

All experiments involving animals at the University of Bern were approved by the Veterinary Service of the Department of Agriculture and Nature of the Canton of Bern (VTHa# BE4/16, BE17/17 and BE127/17). For the QTL mapping study, wild sticklebacks were collected from Nova Scotia, Canada, as previously described [40]. Stickleback care at Stanford University was approved by the Institutional Care and Use Committee (protocol no. 13834).

### Sample collections and genetic crosses

The generation of the *A. quadracus* QTL cross, genotyping markers, and linkage map construction was performed as previously described [40]. The sex of the animals in the QTL cross was determined visually by the presence of red spines in reproductive males and absence of red coloration in females. QTL mapping was performed in R version 4.2.2 using the package R/qtl [83] and a binary model. A total of 380 animals and 269 genotypic markers were used [40]. The bar plot showing phenotypes and genotypes at the top peak marker was generated using R version 4.2.2 and significance of the correlation was assessed using a Chi-square test in R.

Genetic crosses for Pool-seq were made from the following populations, with the wild parents of the crosses collected from: Canal Lake (44.49830, −63.90205) in Nova Scotia (NS), Canada in 2019 by Anna Dalziel; Demarest Lloyd State Park (41.5289936, −70.9833719) in Massachusetts (MA), USA in 2007 by Catherine Peichel; West River Memorial Park (41.314148, −72.956544) in Connecticut (CT), USA in 2009 by Thomas Near (S1 Fig). For each population, a single cross was generated using a single female and a single male. The sex of each F1 offspring was identified by dissection of the gonads, and a fin clip was sampled and preserved in ethanol for DNA extraction and sequencing. For the NS cross, brains were also dissected from 12 males and 12 females from the F1 offspring as well as from the male and female F0 parents used for crossing for further RNA-seq analysis.

Wild populations for whole-genome sequencing of *A. quadracus* were collected from the following localities: Canal Lake (44.49830, −63.90205) in Nova Scotia (NS), Canada in 2021 by Anne Dalziel; Demarest Lloyd State Park (41.5289936, −70.9833719) in Massachusetts (MA), USA in 2007 by Catherine Peichel; and West River Memorial Park (41.314148, −72.956544) in Connecticut (CT), USA in 2021 by Natalie Steinel and Daniel Bolnick. The sex of each individual was identified by dissection of the gonads, and a fin clip was sampled and preserved in 95% ethanol for DNA extraction and sequencing. Total numbers of individuals sequenced for each population and cross are provided in S1 Table.

Note that a previous cytogenetic study of the same MA population used here suggested it had a ZW sex chromosome [7], while a previous cytogenetic study of the same CT population used here did not identify any heteromorphic sex chromosome pair [39].

### DNA and RNA extraction and sequencing

For the male genome assembly, DNA from a single laboratory-reared male from a Canal Lake population cross (Nova Scotia, Canada) was used. For the male assembly, high molecular weight DNA was extracted from the liver following previously described methods [31] and used to prepare a HiFi SMRTbell library for PacBio HiFi sequencing. The blood of the same individual was used to prepare a Hi-C sequencing library using the Phase Genomics Proximo Hi-C animal kit (Phase Genomics, Seattle, WA). One SMRT cell was sequenced on a PacBio Sequel IIe, two SMRT cells were sequenced on a PacBio Revio, and Hi-C libraries were sequenced for 300 cycles on an Illumina NovaSeq S1 flow cell. All library preparation and sequencing were performed by the University of Bern Next Generation Sequencing Platform.

For Pool-seq of F1 offspring of genetic crosses and DNA-sequencing of F0 parents, DNA was extracted by phenol-chloroform extraction, followed by ethanol precipitation. Sequencing libraries were created by standard Illumina DNA TruSeq kits. For the RNA-sequencing of F0 parents and F1 offspring from the NS cross, total brain RNA from the F0 parents and F1 offspring of the NS cross was extracted using Trizol (Life Technologies, Carlsbad, California, USA) following the manufacturer’s instructions. RNA-seq libraries were prepared with the Illumina mRNA TruSeq kit. All libraries were subject to 150bp paired-end sequencing on Illumina NovaSeq SP flow cells by the University of Bern Next Generation Sequencing Platform.

DNA of wild-caught samples for whole-genome sequencing was extracted by phenol-chloroform extraction, followed by ethanol precipitation. Multiplexed haplotagging libraries were prepared as described in [84] with the following modifications in WASH buffer volumes, Tn5 stripping, subsampling and exonuclease reaction. Briefly, DNA were processed in batches of 96 samples. For each sample, 0.75 ng input DNA at 0.15 ng/µl concentration were mixed with 2.5µl haplotagging beads resuspended in 20µl of WASH buffer (20 mM Tris pH8, 50 mM NaCl, 0.1% Triton X-100). We reduced the volume of the tagmentation reaction by using only 5µl of 5x tagmentation buffer (50 mM TAPS pH 8.5 with NaOH, 25 mM MgCl2, 50% N,N-dimethylformamide) and 15 µl of 0.6% SDS for Tn5 stripping following tagmentation. Next, the samples were pooled with 1/3 bead subsampling. This corresponds to a final input DNA of 0.25 ng per sample.

With only 8 pooled samples on the magnetic stand, the buffer was removed, and 20 µl of 1x Lambda Exonuclease buffer, supplemented with 10 units of Exonuclease I (M0293L, New England BioLabs), was added to each sample. Samples were incubated at 48 °C for 20 minutes, and then washed twice for 5 minutes with 150 µl of WASH buffer. DNA library was then amplified using NEBNext® High-Fidelity 2X PCR Master Mix (M0541L, New England BioLabs) in eight 50 µl PCR reaction according to manufacturer’s instructions, using 3 µl of 10 µM TruSeq-F AATGATACGGCGACCACCGAGATCTACAC and TruSeq-R CAAGCAGAAGACGGCATACGAGAT primers, with the following cycling conditions: 10 min at 72°C followed by 30 sec 98°C and 10 cycles of: 98°C for 15 sec, 65°C for 30 sec and 72°C for 60 sec. Libraries were pooled after PCR into a single library pool, size selected using 0.9x volume of Ampure magnetic beads (Beckman Coulter), Qubit quantified, followed by a second size selection with 0.45x and 0.85x volume of Ampure magnetic beads, to remove library longer than 800 bp and smaller than 300 bp, respectively. Pooled libraries were sequenced on a whole S4 lane of Novaseq 6000 (Illumina) instrument with a 151+13+13+151 cycle run setting, such that the run produced 13 and 13 nt in the i7 and i5 index reads, respectively. Sequence data were first converted into fastq format using --create-fastq-for-index-reads using the bcl2fastq program (Illumina). Then we performed beadTag demultiplexing to generate the modified fastq files using a custom demult_fastq program, resulting in a fastq file supplemented with molecular and sample barcode in the header of each read (e.g. BX:Z:A01C02B03D04). This program is available at https://github.com/evolgenomics/haplotagging.

### *Apeltes quadracus* male de novo assembly and annotation

Raw HiC reads were trimmed by Trimmomatic (v 0.36) [85] with a sliding window of 4 bp. The first 13 bp of reads were dropped, and windows of the remaining reads were also dropped with an average quality score below 15. Together with the HiFi reads, two phased assemblies were generated using Hifiasm (0.19.8-r603) with the “Hi-C integration” option and the default parameters [86].

For each haploid assembly, contig scaffolding was conducted using Hi-C proximity guided assembly separately. Trimmed Hi-C reads were first processed with Chromap (v0.2.6) [87] and then assembled by YaHS [88]. After the first round of Hi-C scaffolding, the assembly was revised manually based on the contact map and then scaffolded again. The final step, gap-closing, was run by TGS-GapCloser (1.2.1) [89]. To identify the Y chromosome, we compared two haploid assemblies with the previously published female assembly [41] using mummer 4 [90] and JCVI [91]. The final assembly contained a full set of haploid assembly of autosomes, a haploid X chromosome and a haploid Y chromosome. Assembly quality was evaluated by BUSCO v4 [92,93].

Identification of repeat elements and the establishment of repeat library were conducted by EDTA (2.0.1) [94]. The genome assembly was masked by RepeatMasker (v. 4.1.1) [95]. The RNA-seq data generated from 12 *A. quadracus* males from the NS cross and RNA-seq from NCBI database were used to aid in genome annotation (See S1 Table for details). The raw reads were trimmed by Trimmomatic (v. 0.36) and then mapped against the soft-masked male assembly by Hisat2 (v2.2.1) [96]. Genome annotation was done by EGAPx (v0.2-alpha) with the integration of RNA and protein data of Actinopterygii from NCBI [97]. Lastly, the functional annotation was conducted by eggnog-mapper (v2) [98].

### Short read data processing and SNP calling

All DNA sequencing reads were trimmed by Trimmomatic (v 0.36) [85] with a sliding window of 4 bp. The first 13 bp of all reads were dropped, and windows with an average quality score below 15 were also dropped.

For Pool-seq reads from genetic crosses, trimmed reads were first mapped to the male assembly without the Y chromosome by BWA (v 0.7.11) and sorted with duplicates removed by Picard 2.0.1. Pooplation2 [99] was used to create a sync file containing all the variants for each cross separately.

For linked-reads sequencing from wild populations, trimmed reads were first mapped to the male assembly without the Y chromosome by EMA [100], and remaining unmapped reads were further mapped by BWA (v 0.7.11) [101]. Bam files were sorted, and duplicates were removed by Picard 2.0.1 (http://broadinstitute.github.io/picard). SNP calling was done by GATK 4.1.1 [102]. Vcftools 0.1.16 [103] was used to further filter the SNP matrix with the following criteria: (1) individuals with a mean coverage lower than 6; (2) the population mean depth coverage at the SNP was less than 4x or greater than 40x; (3) the proportion of missing data at the SNP was greater than 0.2 in either the CT population or the NS population; (4) the minor allele frequency of the SNP was less than 0.05. The MA population had poor sequencing quality (likely due to the age of the samples) and was therefore not used for SNP filtering, in order to rescue as much information as possible from this population.

### Identification of the sex determination system and sex chromosome in *A. quadracus*

The sex determination system and sex chromosome were identified in *A. quadracus* using multiple lines of evidence. Using mosdepth 0.3.3 [104] with a sliding window of 20kb and a step size of 10kb, sequencing depth was calculated for both linked read data from wild populations and Pool-seq data from genetic crosses. For the genetic crosses, PoPoolation1 [105] was used for calculating Pi, and PoPoolation2 [99] was used for calculating Fst. For the wild populations, Fst and genetic diversity were calculated by VCFtools 0.1.16 [103] with a sliding window of the same size. In addition, 40 bp male-specific kmers from each population were identified by KmerGO [106], and then mapped against Y assembly to explore the SDR. To further confirm the sex determination pattern, RNA-seq data from the parents and offspring of the NS cross was fed into read2snp 2.0 [107] to obtain SNP data, and also fed into Trinity 2.11.2 [108] to obtain a de novo transcriptome assembly. The output SNP array and assembly were then processed by SEX-DETector [42].

### Identification of population-specific inversions

To determine whether there are inversions on the sex chromosome, three methods were used. First, the linked-read sequences were used to identify shared barcodes among any pairs of windows of 10kb on each chromosome. Windows with shared barcodes were divided into two categories: windows that are adjacent, and windows that are 500kb apart on the same chromosome. Putative inversions were identified based on the number of shared barcodes between 500 kb apart non-adjacent window pairs. Second, inversions on the sex chromosomes were identified by LEVIATHAN V1.0.2 [109]. Third, screening of bam files for split and discordantly mapped read pairs near the breakpoints of inversion was done by IGV 2.14.1 [110]. Last, genotypes of inversions were determined by the divergence and diversity pattern between sexes within the inversion as well as the SNP density plot generated by VCFtools 0.1.16 at the individual level. The above analyses were not conducted in the MA population due to the poor sequencing quality.

Further, we used the MCScan in JCVI package [111] to compare synteny among stickleback species and *A. flavidus* on the gene level in order to determine the ancestry of the population-specific inversions.

### Pattern of molecular evolution within inversions

Molecular evolution on the sex chromosome was analyzed in genes present on both X and Y chromosomes. Single-copy orthologues were identified using Blast (2.14.1) [112] and filtered following reciprocal blast hits. Mapping of gene pairs on the sex chromosome was first conducted by PRANK [113] and then filtered by Gblocks (0.91b) [114] to exclude non-conserved regions. dN, dS and dN/dS ratio between the X and Y alleles were further calculated by the CodeML module in PAML 4.9 [115].

Degeneration of the Y chromosome usually appears in two forms: (1) the loss of genes either due to complete deletion or degeneration such that the Y allele can no longer be aligned to the X allele; or (2) accumulation of loss-of-function mutations. To identify the genes that degenerated on the *A. quadracus* sex chromosomes, we calculated male to female read-depth ratios of each gene for each wild population by mosdepth. Further, loss-of-function mutations were identified by snpEff v5.1 [116] separately for each sex and each population. Fixed loss-of-function mutations were identified with an allele frequency greater than 0.9 in females or 0.45 in males. The above analyses were not conducted on the MA population.

### Identification of potential sex determination genes

Because the fourspine stickleback genome assembly was obtained from a male individual, genes on X and Y chromosomes can be directly compared with each other. Therefore, we examined the functions of genes on the Y chromosome that have no BLAST hiton autosomes or the X chromosome, and genes within the SDR that have one-to-one orthologs between the X and Y chromosomes by blasting them against the NCBI nr database. For gene pairs with nonsynonymous changes between X and Y chromosomes, SIFT [43] was applied to predict the effects of amino acid changes. In addition, published candidate sex determination genes of teleost fish [59] were blasted against our male assembly to search for homologues that mapped to the sex chromosome pair.

## Supporting information

File containing 11 supplementary figures.

## Data availability

All sequences are uploaded and available on the NCBI Sequence Read Archive (S1 Table). A link to the sequences for review is available at: https://eur03.safelinks.protection.outlook.com/?url=https%3A%2F%2Fdataview.ncbi.nlm.nih.gov%2Fobject%2FPRJNA1164644%3Freviewer%3Dqs1lqc1d5ug414kr516isjdtsr&data=05%7C02%7Czuyao.liu%40unibe.ch%7C84100b64b760459429ea08dce6c3a8b7%7Cd400387a212f43eaac7f77aa12d7977e%7C1%7C0%7C638638975242759207%7CUnknown%7CTWFpbGZsb3d8eyJWIjoiMC4wLjAwMDAiLCJQIjoiV2luMzIiLCJBTiI6Ik1haWwiLCJXVCI6Mn0%3D%7C0%7C%7C%7C&sdata=4mNSrwDd2toO39VClXH%2B9enqZmdqUGmqjvcz%2BaxdjWg%3D&reserved=0

## Acknowledgments

We thank Daniel Jeffries and Mark Kirkpatrick for comments on the manuscript, Daniel Bolnick, Anne Dalziel, Thomas Near, and Natalie Steinel for collecting wild fish, Melanie Hiltbrunner, Shaugnessy McCann, and Verena Saladin for making crosses, Melanie Hiltbrunner and Nicole Nesvadba for performing extractions, and the University of Bern Next Generation Sequencing Platform for library preparation and sequencing.

## Supporting Information

**S1 Fig. Fourspine stickleback sampling locations in this study**. Red dots represent sampled populations. Connecticut (CT), USA; Massachusetts (MA), USA; Nova Scotia (NS), Canada; Samples used in the QTL analysis are labeled with brackets.

**S2 Fig. Male-to-female depth ratio across the genome with pool-seq data from genetic crosses of three populations (CT, NS, and MA).** Raw depth values were normalized to eliminate the difference between two sexes. The size of the sliding window is 20kb and the step size is 10kb. Chromosomes are indicated on the X-axis, and the normalized depth ratio is shown on the Y-axis. Dark and light blue regions indicate the different chromosomes.

**S3 Fig. Genomic distributions of normalized male-to-female depth ratio in 20kb sliding windows calculated from linked-read data from the three populations (CT, NS, and MA).** Alternating colors in each panel are used to highlight the different chromosomes. Red and oranges lines represent the smoothed values.

**S4 Fig. Genetic differentiation (Fst) between males and females from pool-seq data from three crosses (CT, NS, and MA).** The size of the sliding window is 20kb and the step size is 10kb. Chromosomes are indicated on the X-axis, and the Fst values are shown on the Y-axis. Purple and yellow regions indicate the different chromosomes.

**S5 Fig. Genetic differentiation (Fst) between males and females on chromosome 23 calculated from pool-seq data from three genetic crosses**. The Connecticut (CT) cross is in coral, the Massachusetts (MA) cross is in yellow, and the Nova Scotia (NS) cross is in light blue. The locations of the inversions are also indicated. SDR is shown in the grey box. Note that all sequences are aligned to the X chromosome assembly.

**S6 Fig. Genomic distribution of genetic diversity (Pi) in 20kb sliding windows within males and females calculated from pool-seq data from three genetic crosses (CT, NS, and MA).** Chromosomes are indicated on the X-axis, and the values of genetic diversity (Pi) are shown on the Y-axis. Red dots represent males and cyan dots represent females.

**S7 Fig. Distribution of genetic diversity (Pi) within males and females in 20kb sliding windows on chromosome 23 calculated from pool-seq data from the three genetic crosses (CT, NS, and MA).** The position on the X chromosome is given on the X-axis, and the values of genetic diversity (Pi) are shown on the Y-axis. Red dots represent males and cyan dots represent females. Note that all sequences are aligned to the X chromosome assembly.

**S8 Fig. Genomic distribution of genetic diversity (Pi) in 20kb sliding windows within males and females calculated from linked-read data from three populations (CT, NS, and MA).** Chromosomes are indicated on the X-axis, and the values of genetic diversity (Pi) are shown on the Y-axis. Red dots represent males and cyan dots represent females.

**S9 Fig. Segregation patterns of inversions in genetic crosses inferred from pool-seq data.** Red bars represent the X-specific inversion, and blue bars represent the Y-specific inversion. For the MA cross, the number of individuals with each genotype is assumed to be equal within a sex.

S10 Fig. Synteny map of the X chromosome and Y chromosome assemblies generated from the NS population of *A. quadracus* with homologous chromosomes from two other stickleback species (*G. aculeatus*, *P. pungitius*) and an outgroup species (*A. flavidus*). Blue and green bars represent genes. Grey lines are syntenic blocks between species.

**S11 Fig. Distributions of the density of male-specific kmers (40bp) on the Y chromosome.** Male-specific kmers were calculated using linked-reads sequences from three wild populations from Connecticut (CT), Nova Scotia (NS), and Massachusetts (MA). A sliding window of 20kb was used

**S1 Table. Sample information and accession numbers for sequencing data in this study.**

**S2 Table. SEX-Detector results.** The first column represents the model used in each run. The second column shows the number of sex-linked transcripts detected.

**S3 Table. Identification of inversions in the CT and NS populations.** Evidence comes from LEVIATHAN results, split reads, and read pairs with long insertion sizes.

**S4 Table.** Genes with loss-of-function mutations on X and Y chromosomes.

**S5 Table. Y-specific genes, genes with one-to-one homologues on the X and Y chromosomes within the sex determination region, and genes around 12Mb.** Genes of interest are labeled with bold font.

**S1 Appendix. Plots of individual heterozygosity from CT, NS and MA populations.**

