## Supplementary material for "The fourspine stickleback (*Apeltes quadracus*) has an XY sex chromosome system with polymorphic inversions on both X and Y chromosomes": File containing 11 supplementary figures.

### Supporting Information

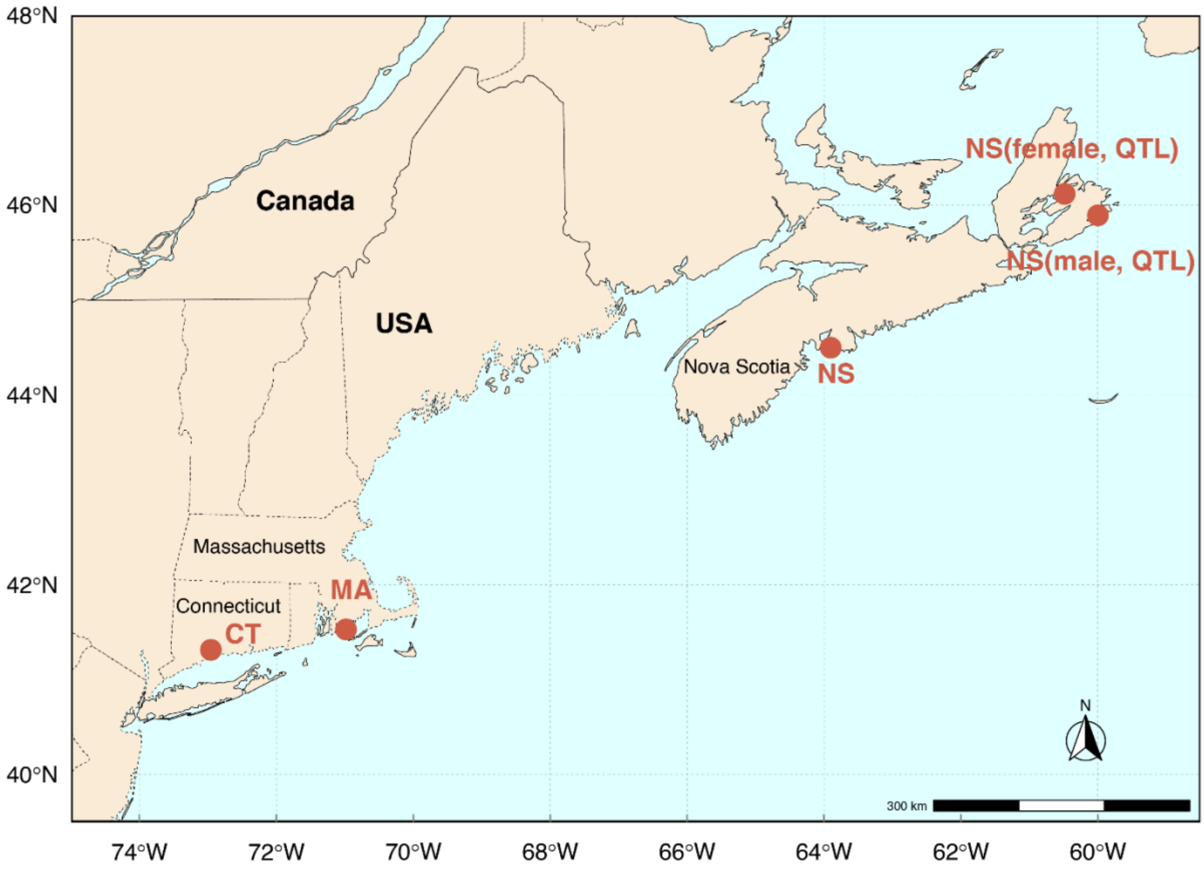


**S1 Fig. Fourspine stickleback sampling locations in this study**. Red dots represent sampled populations. Connecticut (CT), USA; Massachusetts (MA), USA; Nova Scotia (NS), Canada; Samples used in the QTL analysis are labeled with brackets.

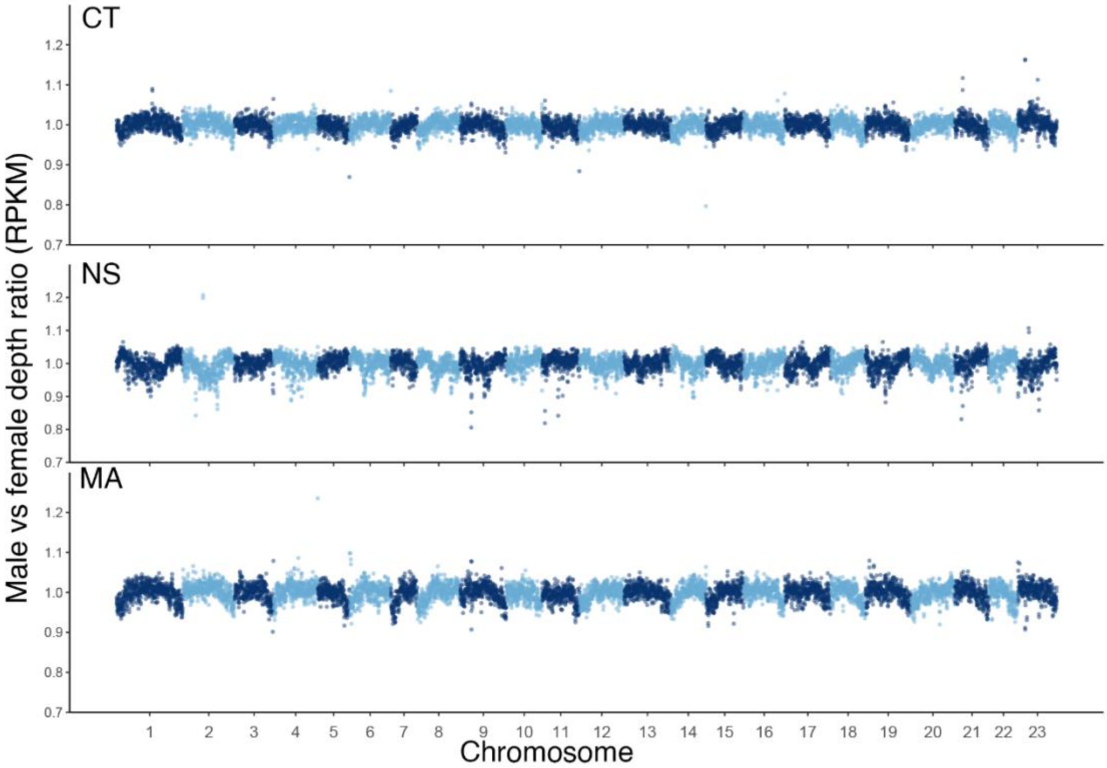


**S2 Fig. Male-to-female depth ratio across the genome with pool-seq data from genetic crosses of three populations (CT, NS, and MA).** Raw depth values were normalized to eliminate the difference between two sexes. The size of the sliding window is 20kb and the step size is 10kb. Chromosomes are indicated on the X-axis, and the normalized depth ratio is shown on the Y-axis. Dark and light blue regions indicate the different chromosomes.

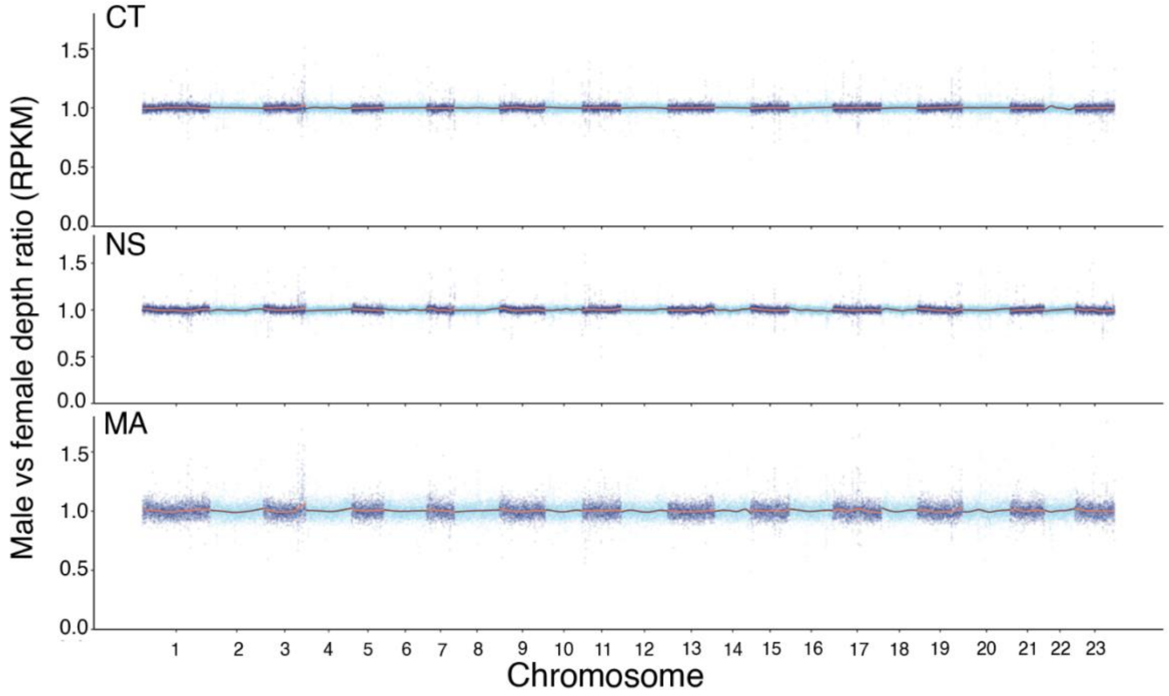


**S3 Fig. Genomic distributions of normalized male-to-female depth ratio in 20kb sliding windows calculated from linked-read data from the three populations (CT, NS, and MA).** Alternating colors in each panel are used to highlight the different chromosomes. Red and oranges lines represent the smoothed values.

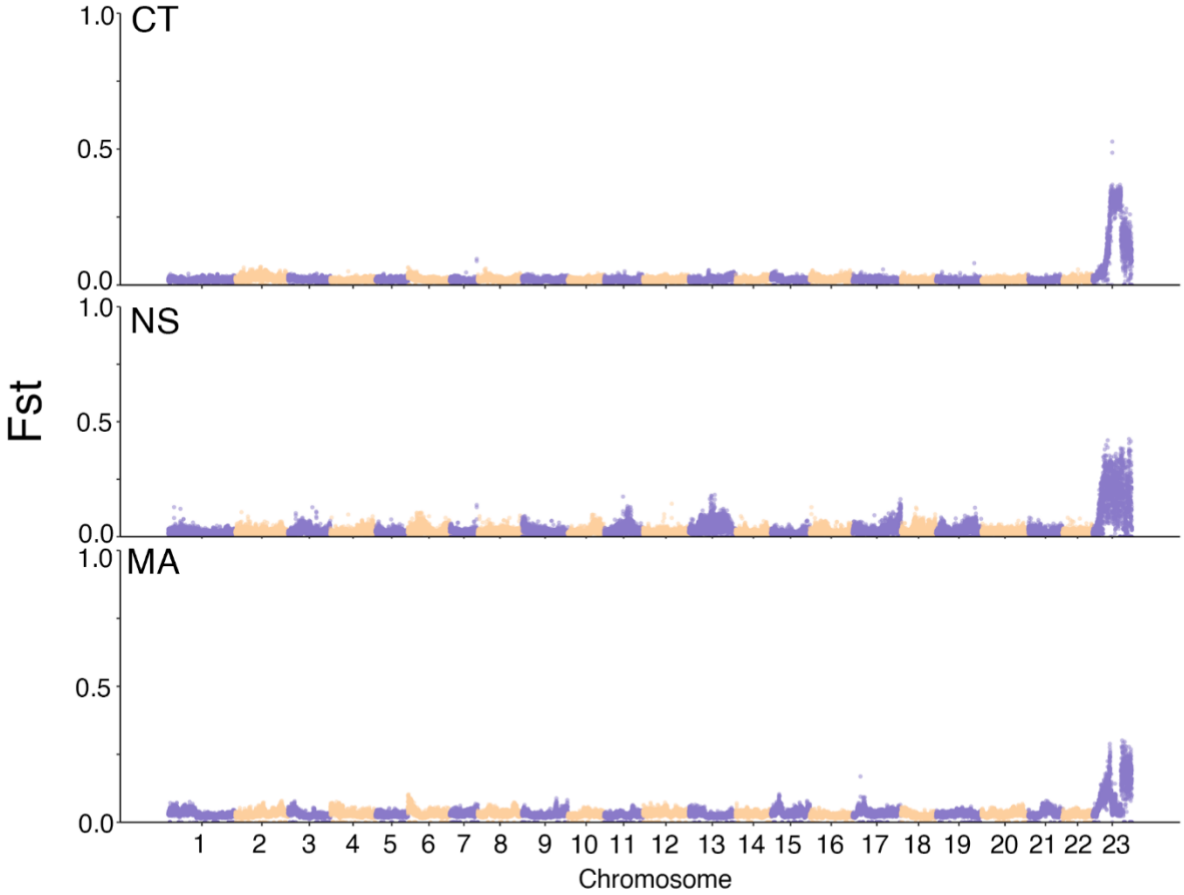


**S4 Fig. Genetic differentiation (Fst) between males and females from pool-seq data from three crosses (CT, NS, and MA).** The size of the sliding window is 20kb and the step size is 10kb. Chromosomes are indicated on the X-axis, and the Fst values are shown on the Y-axis. Purple and yellow regions indicate the different chromosomes.

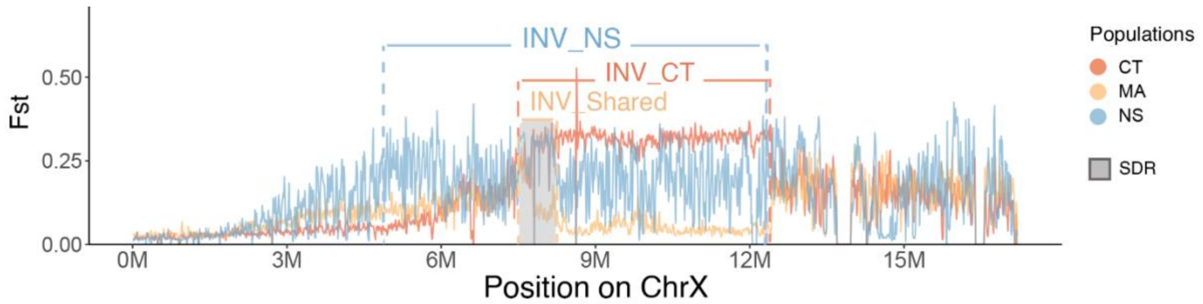


**S5 Fig. Genetic differentiation (Fst) between males and females on chromosome 23 calculated from pool-seq data from three genetic crosses**. The Connecticut (CT) cross is in coral, the Massachusetts (MA) cross is in yellow, and the Nova Scotia (NS) cross is in light blue. The locations of the inversions are also indicated. SDR is shown in the grey box. Note that all sequences are aligned to the X chromosome assembly.

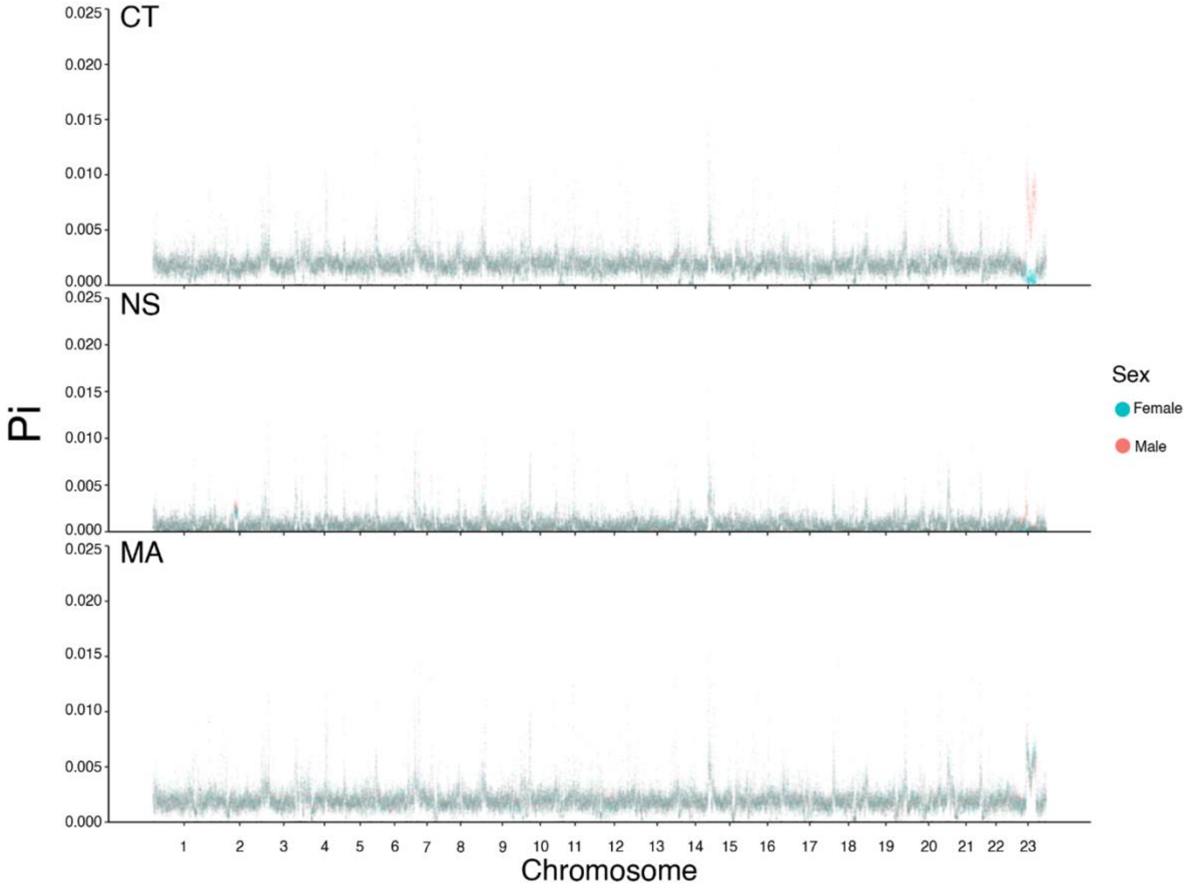


**S6 Fig. Genomic distribution of genetic diversity (Pi) in 20kb sliding windows within males and females calculated from pool-seq data from three genetic crosses (CT, NS, and MA).** Chromosomes are indicated on the X-axis, and the values of genetic diversity (Pi) are shown on the Y-axis. Red dots represent males and cyan dots represent females.

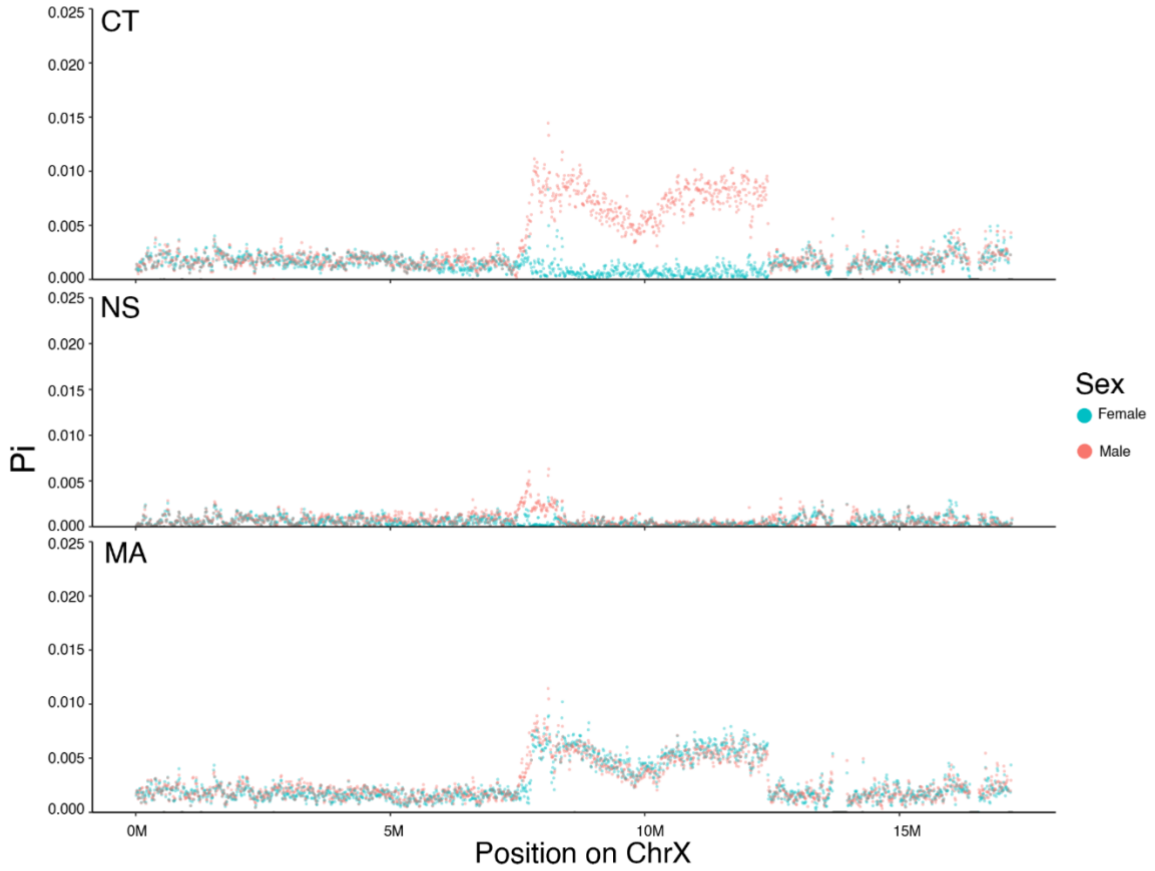


**S7 Fig. Distribution of genetic diversity (Pi) within males and females in 20kb sliding windows on chromosome 23 calculated from pool-seq data from the three genetic crosses (CT, NS, and MA).** The position on the X chromosome is given on the X-axis, and the values of genetic diversity (Pi) are shown on the Y-axis. Red dots represent males and cyan dots represent females. Note that all sequences are aligned to the X chromosome assembly.

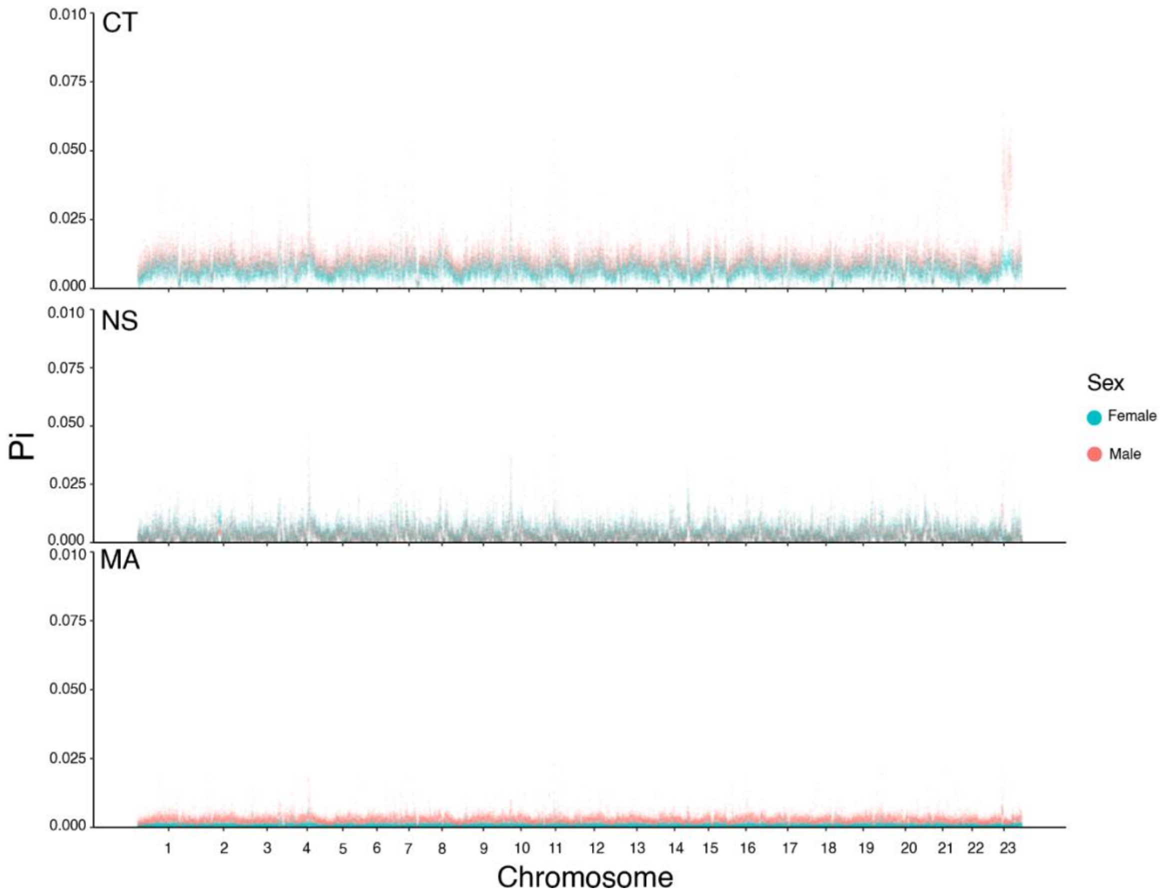


**S8 Fig. Genomic distribution of genetic diversity (Pi) in 20kb sliding windows within males and females calculated from linked-read data from three populations (CT, NS, and MA).** Chromosomes are indicated on the X-axis, and the values of genetic diversity (Pi) are shown on the Y-axis. Red dots represent males and cyan dots represent females.

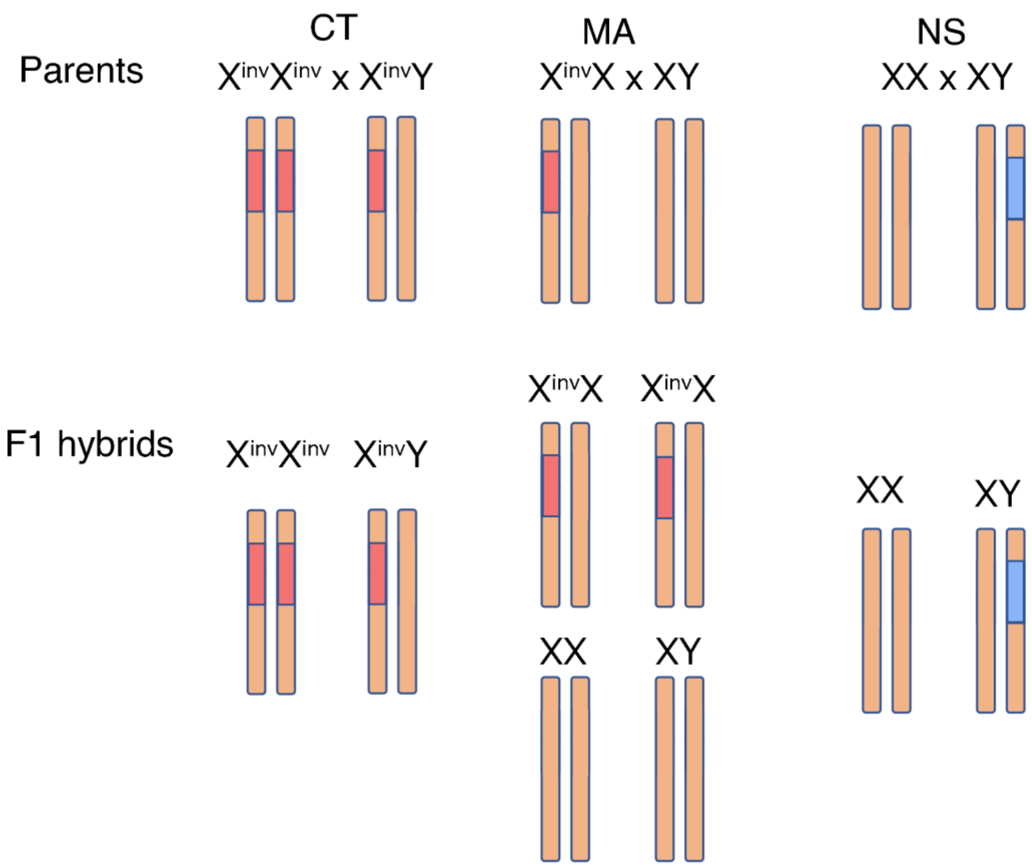


**S9 Fig. Segregation patterns of inversions in genetic crosses inferred from pool-seq data.** Red bars represent the X-specific inversion, and blue bars represent the Y-specific inversion. For the MA cross, the number of individuals with each genotype is assumed to be equal within a sex.

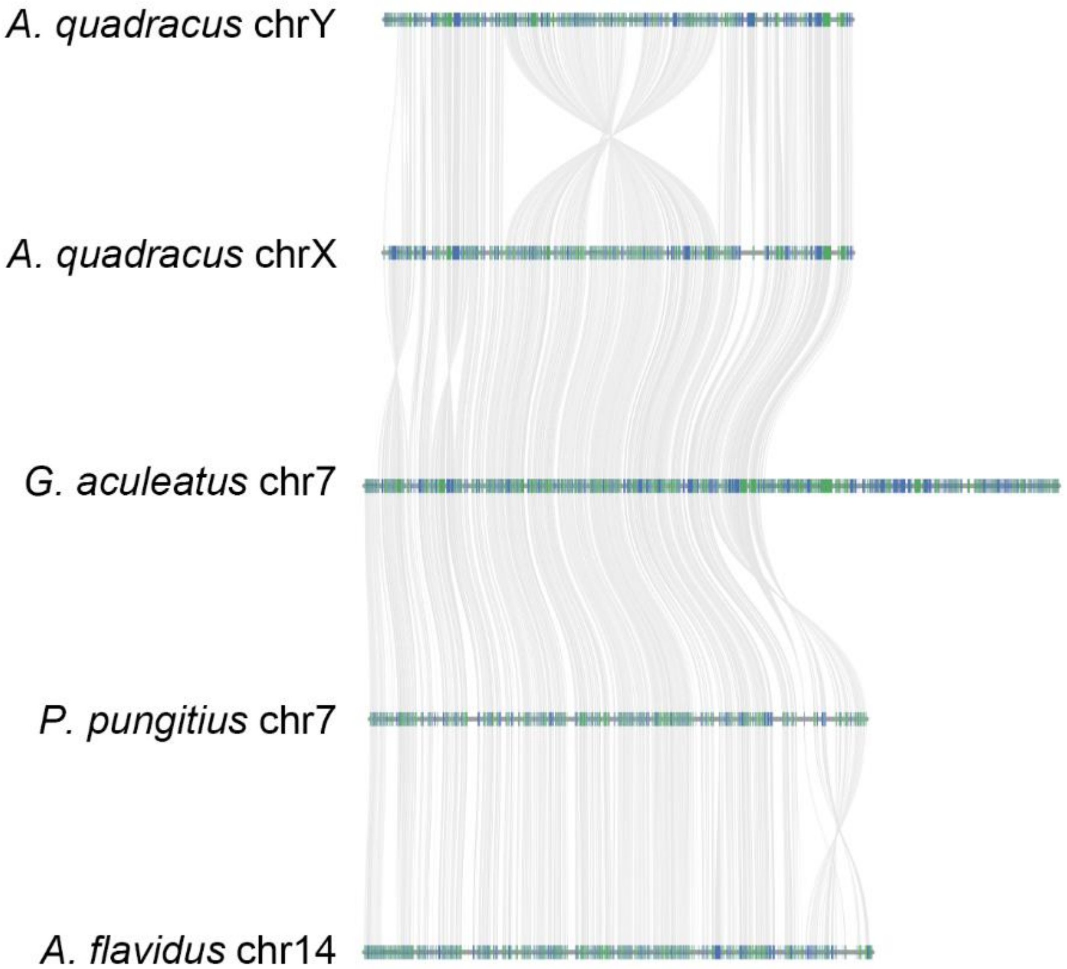


**S10 Fig. Synteny map of the X chromosome and Y chromosome assemblies generated from the NS population of *A. quadracus* with homologous chromosomes from two other stickleback species (*G. aculeatus*, *P. pungitius*) and an outgroup species (*A. flavidus*).** Blue and green bars represent genes. Grey lines are syntenic blocks between species.

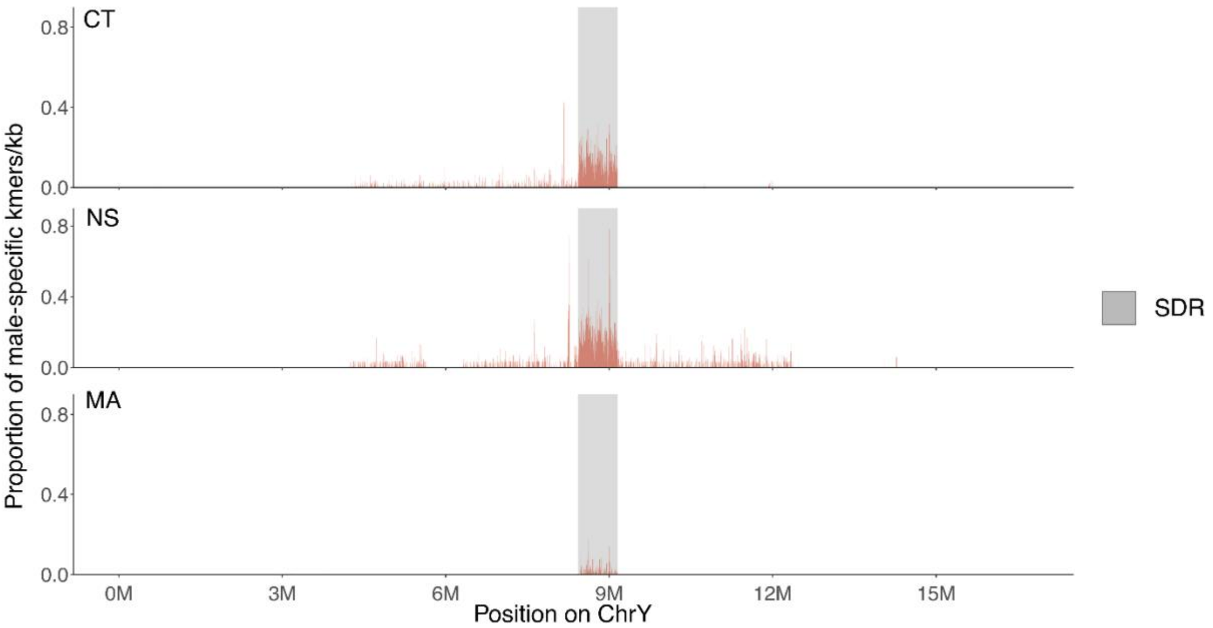


**S11 Fig. Distributions of the density of male-specific kmers (40bp) on the Y chromosome.** Male-specific kmers were calculated using linked-reads sequences from three wild populations from Connecticut (CT), Nova Scotia (NS), and Massachusetts (MA). A sliding window of 20kb was used.

#### Tables

**S1 Table. Sample information and accession numbers for sequencing data in this study.**

**S2 Table. SEX-Detector results.** The first column represents the model used in each run. The second column shows the number of sex-linked transcripts detected.

**S3 Table. Identification of inversions in the CT and NS populations.** Evidence comes from LEVIATHAN results, split reads, and read pairs with long insertion sizes.

**S4 Table.** Genes with loss-of-function mutations on X and Y chromosomes.

**S5 Table. Y-specific genes, genes with one-to-one homologues on the X and Y chromosomes within the sex determination region, and genes around 12Mb.** Genes of interest are labeled with bold font.

#### Appendix

**S1 Appendix. Plots of individual heterozygosity from CT, NS and MA populations.**
